# Linked-read sequencing enables haplotype-resolved resequencing at population scale

**DOI:** 10.1101/2020.01.15.907261

**Authors:** Dave Lutgen, Raphael Ritter, Remi-André Olsen, Holger Schielzeth, Joel Gruselius, Phil Ewels, Jesús T. García, Hadoram Shirihai, Manuel Schweizer, Alexander Suh, Reto Burri

## Abstract

The feasibility to sequence entire genomes of virtually any organism provides unprecedented insights into the evolutionary history of populations and species. Nevertheless, many population genomic inferences – including the quantification and dating of admixture, introgression and demographic events, and the inference of selective sweeps – are still limited by the lack of high-quality haplotype information. In this respect, the newest generation of sequencing technology now promises significant progress. To establish the feasibility of haplotype-resolved genome resequencing at population scale, we investigated properties of linked-read sequencing data of songbirds of the genus *Oenanthe* across a range of sequencing depths. Our results based on the comparison of downsampled (25x, 20x, 15x, 10x, 7x, and 5x) with high-coverage data (46-68x) of seven bird genomes suggest that phasing contiguities and accuracies adequate for most population genomic analyses can be reached already with moderate sequencing effort. At 15x coverage, phased haplotypes span about 90% of the genome assembly, with 50 and 90 percent of the phased sequence located in phase blocks longer than 1.25-4.6 Mb (N50) and 0.27-0.72 Mb (N90), respectively. Phasing accuracy reaches beyond 99% starting from 15x coverage. Higher coverages yielded higher contiguities (up to about 7 Mb/1Mb (N50/N90) at 25x coverage), but only marginally improved phasing accuracy. Finally, phasing contiguity improved with input DNA molecule length; thus, higher-quality DNA may help keeping sequencing costs at bay. In conclusion, even for organisms with gigabase-sized genomes like birds, linked-read sequencing at moderate depth opens an affordable avenue towards haplotype-resolved genome resequencing data at population scale.

## Introduction

The possibility to sequence entire genomes at population-scale has nothing short of revolutionized molecular ecology. Genome-wide data are now routinely used to infer populations’ histories of demography, admixture, and introgression (Beichman *et al.* 2018; Gompert & Buerkle 2013; Taylor & Larson 2019); investigations of genetic diversity along genomes have provided unprecedented insights into the distribution of most differentiated regions across genomes (e.g. Burri *et al.* 2015; Jones *et al.* 2012; Martin *et al.* 2013; Soria-Carrasco *et al.* 2014) and the processes driving their evolution (reviewed e.g. in Burri 2017; Martin & Jiggins 2017; Ravinet *et al.* 2017); and numerous phenotypes have been mapped to genomic regions underpinning their expression (e.g. Kupper *et al.* 2016; Lamichhaney *et al.* 2015; Santure & Garant 2018; Schielzeth *et al.* 2018). Enabling these insights, studies of genome-wide variation continue contributing to our understanding of the evolutionary histories of phenotypes, populations, and species at an ever increasing pace.

Nonetheless, deeper insights into the molecular ecology of natural populations – in particular the extent and timing of admixture and introgression, and the distribution of genomic regions that underwent selective sweeps – can be achieved by the integration of yet usually difficult to obtain haplotype information, that is, information on the gametic phase of genetic variants (e.g. Palamara *et al.* 2012). Haplotype structure reflects some of the most significant footprints left by admixture (Buerkle & Rieseberg 2008; Fisher 1949, 1954; Pool & Nielsen 2009), selective sweeps (e.g. Sabeti *et al.* 2002; Voight *et al.* 2006), or a combination thereof (e.g. Shchur *et al.* 2019). Haplotype tracts that enter populations (according to context referred to as ‘migrant tracts’ or ‘ancestry tracts’) are progressively broken down to smaller size as recombination events accumulate over evolutionary time (e.g. Janzen *et al.* 2018; Pool & Nielsen 2009). According to this recombination clock (Moorjani *et al.* 2016), long haplotype tracts are of more recent origin than shorter ones. Thus, migrant tracts that entered populations/species through admixture/ introgression recently are expected to be longer than migrant tracts of older origin (Pool & Nielsen 2009). The size distribution of migrant tracts thus enables powerful insights into histories of admixture and introgression (e.g. Palamara & Pe’er 2013). Likewise, long haplotype tracts that reached high frequency in a population must have done so in a timeframe that did not allow recombination to break them down – thus fast and likely by positive selection (e.g. Sabeti *et al.* 2002; Voight *et al.* 2006). Therefore, whereas current inference methods that predominantly rely on allelic states and/or allele frequencies may be limited in their power to distinguish among complex demographic hypotheses and detecting selective sweeps, these tasks may be greatly facilitated by haplotype-based approaches (Gompert & Buerkle 2013).

However, the application of haplotype-based approaches to natural populations has yet been stalled by the difficulty to obtain phased genomic data, that is, data for which the gametic phase of genetic variants is known. Although gametic phase can be inferred statistically (e.g. Delaneau *et al.* 2012; Stephens *et al.* 2001), this requires numbers of individuals that surmount what sequencing experiments in natural populations can usually achieve. With a limited number of individuals, statistical phasing has low accuracy and reduced power (Browning & Browning 2011; Choi *et al.* 2018), and may limit for instance the inference of selective sweeps (Nadachowska-Brzyska *et al.* 2019). Therefore, haplotype-based approaches to infer admixture and selection have largely been limited to humans, where thousands of available genome sequences enable statistical phasing with reasonable confidence (e.g. Loh *et al.* 2016; O’Connell *et al.* 2016). A notable exception is a recent study on natural populations of sea bass that used parent-offspring trios to accurately infer the gametic phases of genome-wide data (Duranton *et al.* 2018). However, this approach is expensive, laborious, and not applicable to all species. Alternative approaches to obtain phased genomic data are thus called for.

The latest generation of genome sequencing technologies now offers promising avenues towards cost-efficient haplotype-resolved genome sequencing – sequencing that directly determines gametic phase from template DNA molecules (Snyder *et al.* 2015). Long-read technologies, such as offered by Pacific Biosciences and Oxford Nanopore, determine genome sequences from single molecules. These approaches yield sequences with lengths up to tens of kilobases (kb), and accordingly provide phase information across at least the same scale. Still, both methods require high amounts of input DNA and remain prohibitively expensive for population-scale sequencing experiments. On the other hand, affordable standard short-read sequencing by itself provides phasing over at most a couple of hundred base pairs. However, dedicated ‘linked-read’ library preparation protocols now enable haplotype-resolved sequencing based on short-read sequencing (Chen *et al.* 2019; Redin *et al.* 2019; Snyder *et al.* 2015) that outperforms statistical phasing of human data (Choi *et al.* 2018). One approach to the preparation of such linked-read performs library preparation in emulsion droplets that each contain a very limited number of DNA molecules. The addition of droplet-specific barcodes during library preparation distinguishes short reads from different DNA molecules and thereby enables efficient resolution of haplotypes of lengths up to over hundred kb (Weisenfeld *et al.* 2017; Zheng *et al.* 2016). Alternatively, transposases are used to introduce barcodes that distinguish sequence fragments originating from different DNA molecules (Chen *et al.* 2019). Thus, while for the time being long-read sequencing of multiple individuals remains prohibitively expensive (but see Weissensteiner *et al.* 2019), on top of enabling cost-efficient and contiguous *de novo* genome assemblies of gigabase-sized genomes (e.g. Boman *et al.* 2019; Kinsella *et al.* 2019; Schweizer *et al.* 2019a; Toomey *et al.* 2018), linked-read sequencing opens the scope for haplotype-resolved sequencing at population scale.

Here, we set out to determine the linked-read sequencing effort required to obtain accurate and long-range phase information. Such information can help maximizing the cost-efficiency of linked-read sequencing for haplotype-resolved sequencing at population scale. To this end, we performed a downsampling experiment on seven ca 1 gigabase-sized bird genomes originally sequenced at 48-68x coverages, and determined phasing contiguities and accuracies at coverages of 25x down to 5x. Our results suggest that adequate phase block contiguities and phasing accuracies can be reached with read-depths as low as 15x. This implies that haplotype-resolved sequencing is feasible at population scale, and opens new perspectives for the implementation of high-quality phase information in demographic reconstructions (Harris & Nielsen 2013; Lawson *et al.* 2012; Palamara & Pe’er 2013), genome scans for selective sweeps and reproductive incompatibilities (Ferrer-Admetlla *et al.* 2014; Sabeti *et al.* 2002; Sedghifar *et al.* 2016; Tang *et al.* 2007; Voight *et al.* 2006), QTL mapping (Schielzeth & Husby 2014) and conservation genomics (Duranton *et al.* 2019; Leitwein *et al.*).

## Methods

### DNA extraction, library preparation, and genome sequencing

We sequenced the genomes of seven individuals of the four species in the *Oenanthe hispanica-pleschanka-melanoleuca-cypriaca* complex (Schweizer *et al.* 2019a) within the open-habitat chats (Aliabadian *et al.* 2012; Schweizer *et al.* 2019b) (**Table 1**). To this end, DNA was extracted from blood or tissue conserved in ethanol from each two western black-eared wheatears (*O. hispanica*), two pied wheatears (*O. pleschanka*), two eastern black-eared wheatears (*O. melanoleuca*), and a Cyprus wheatear (*O. cypriaca*) (**Table 1**) using the MagAttract HMW DNA kit (Qiagen, Hilden, Germany). Digestion was performed in 360 ul of buffer ATL, 440 ul of buffer AL, and 20 ul of proteinase K starting at 56°C for one hour and 37°C overnight. Additional 20 ul of proteinase K were added in the morning, and digestion completed at 56°C. Total time of digestion was about 16 hours. DNA extraction then followed the manufacturer’s recommendations.

**Table 1.** Sample information.

| Individual | Species | (Museum) No.* | Year | Tissue | Coverage [x] | Mol. Length [kb]** |
| --- | --- | --- | --- | --- | --- | --- |
| 7359_104 | *O. melanoleuca* | YPM 101348 | 2005 | Muscle | 68 | 10.6 |
| 8854_101 | *O. pleschanka* | MCZ 349924 | 2012 | Muscle | 54 | 17.9 |
| 8854_102 | *O. melanoleuca* | A1167 | 2000 | Blood | 54 | 47.8 |
| 8854_103 | *O. hispanica* | 2917 | 2009 | Blood | 52 | 33.1 |
| 8854_104 | *O. hispanica* | 2919 | 2009 | Blood | 56 | 58.9 |
| 8854_105 | *O. pleschanka* | 16 | 2012 | Blood | 46 | 37.3 |
| 8854_106 | *O. cypriaca* | 19 | 2012 | Blood | 60 | 63.8 |

\*Individual identifiers and where applicable museum numbers are provide. MCZ, Museum of Comparative Zoology Harvard; YPM, Yale Peabody Museum
\*\* Mean input DNA molecule length as determined by Supernova 2.1

Linked-read sequencing libraries were prepared using Chromium Genome library kits (10X Genomics) and each library was sequenced on half a lane of an Illumina HiSeq X flowcell.

### Reference genome draft assembly and curation

Here, we assembled the genome of the Cyprus wheatear (*O. cypriaca*; **Table 1**) using the 10X Genomics Supernova 2.1 assembler (a significantly less contiguous Supernova 1 assembly of O. melanoleuca YPM 101348 was published in Schweizer *et al.* 2019a and is available from Dryad doi:10.5061/dryad.6d006j3). This resulted in a merged, pseudohaploid assembly of 1.08 Gb length with a scaffold N50 of 25.15 Mb in 23,105 scaffolds. To assess assembly completeness, we evaluated the presence, completeness, and copy number of avian benchmarking universal single-copy orthologs (BUSCOs, aves_odb9, creation date: 2016/02/13) as assessed by BUSCO version 3 (Simão *et al.* 2015). Of 4,915 BUSCOS, 90.8% (4,462) were complete and single copy, 1.3% (62) were complete and duplicated, 4.4% (214) were fragmented, and 3.6% (177) were missing.

To remove duplicate scaffolds of at least 99% identity, we ran the dedupe procedure in BBTools (https://sourceforge.net/projects/bbmap/) allowing up to 7,000 edits. This reduced the assembly to 11,030 scaffolds. We then aimed to ensure that all duplicate scaffolds were removed and retain only scaffolds whose integrity can be confirmed by the presence of syntenic regions in another songbird genome. To this end, we performed a lastz alignment against the collared flycatcher assembly version 1.5 (Kawakami *et al.* 2014), which is the highest-quality assembly available from the Muscicapidae family. For this we used lastz 1.04 (Harris 2007) with settings M=254, K=4500, L=3000, Y=15000, C=2, T=2, and --matchcount=10000. This resulted in 295 scaffolds with unique hits in the flycatcher assembly. The final assembly spanned 970 Mb, with a scaffold N50 of 32.13 Mb. All following analyses used this Cyprus wheatear draft genome assembly as reference.

### Evaluation of phasing accuracy, genotyping accuracy, and phase set contiguity

To obtain proxies for phasing and genotyping accuracy, we assumed the phasing and genotyping of the full data for each individual (46-68x coverage, **Table 1**) to be correct. Mismatches in phase and genotype of the downsampled with the full data were considered as inaccurate phasing and genotyping. To evaluate these accuracies at different coverages, we downsampled the full data to 25x, 20x, 15x, 10x, 7x, and 5x coverage directly in the LongRanger pipeline, which was also used for SNP calling and genotyping, and phasing (Longranger in GATK mode with GATK version 3.8.0, McKenna *et al.* 2010). One-fold genome size for the downsampling procedure was assumed 1.08 Gb, which corresponds to the Supernova version 2.1 merged/pseudohaploid assembly length.

To determine whether phasing corresponded between the downsampled and full data sets, we parsed the genotypes including phasing information for SNPs that fulfilled the following criteria: 1) biallelic, heterozygous SNPs, which 2) passed LongRanger filter criteria, and 3) for which the genotype was identical between the downsampled and full data set. The number and length of phase sets differ between the full and the downsampled data set. Moreover, the order in which maternal or paternal haplotypes are output can also differ between the full and downsampled data and needs to be matched before further analysis. To this end, before estimating phasing mismatches, we determined all combinations of phase sets between the full and downsampled data. For each phase set combination we then determined whether the phasing was resolved in the same order in the two data sets or whether it had to be inversed for the phasing to be compared. For phase set combinations for which the majority of phasings were wrong if retained in the original order (as indicated by a majority of inversed phases), we inversed the phasing order before determining phasing accuracy.

Genotyping accuracies were inferred as the percentage of SNPs with matching genotype between the downsampled and full data sets (before applying the three criteria above).

Phasing contiguity was estimated in terms of phase set N50 and N90. These values were estimated in reference to the combined lengths of all phase sets. That is, phase sets equal and longer then N50/N90 cover 50/90 percent of the number of base pairs contained in all phase sets combined. Furthermore, we estimated the proportion of the genome that was phased.

### Statistical analysis

To determine the effect of sequencing coverage, SNP filtering and input DNA molecule length on phasing and genotyping accuracy and on phasing contiguity, we ran linear models in R 3.5.2. N50 and N90 were log-transformed, accuracies transformed as -log(100-x), and explanatory variables centred. We determined input DNA molecule length for each sample in Supernova version 2.1.

## Results

### Phasing contiguity

Phasing contiguity increased by one to two orders of magnitude with increasing sequencing coverage, and was significantly higher for longer input DNA molecule length (**Figure 1, Table 2**).

**Table 2.** Effect of coverage and molecule length on phasing contiguity (N50, multiple R^2^=0.91; N90, multiple R^2^=0.88; 206 degrees of freedom).

|  | N50 |  |  | N90 |  |  |
| --- | --- | --- | --- | --- | --- | --- |
|  | Estimate | t | p | Estimate | t | p |
| Intercept | 13.482 | 427.234 | $< 10^{-3}$ | 11.907 | 363.976 | $< 10^{-3}$ |
| Coverage | 0.181 | 40.725 | $< 10^{-3}$ | 0.144 | 31.288 | $< 10^{-3}$ |
| Molecule Length | 0.036 | 21.014 | $< 10^{-3}$ | 0.037 | 21.041 | $< 10^{-3}$ |
| Coverage x Molecule Length | -0.001 | -5.771 | $< 10^{-3}$ | -0.001 | -5.947 | $< 10^{-3}$ |

**Figure 1.**
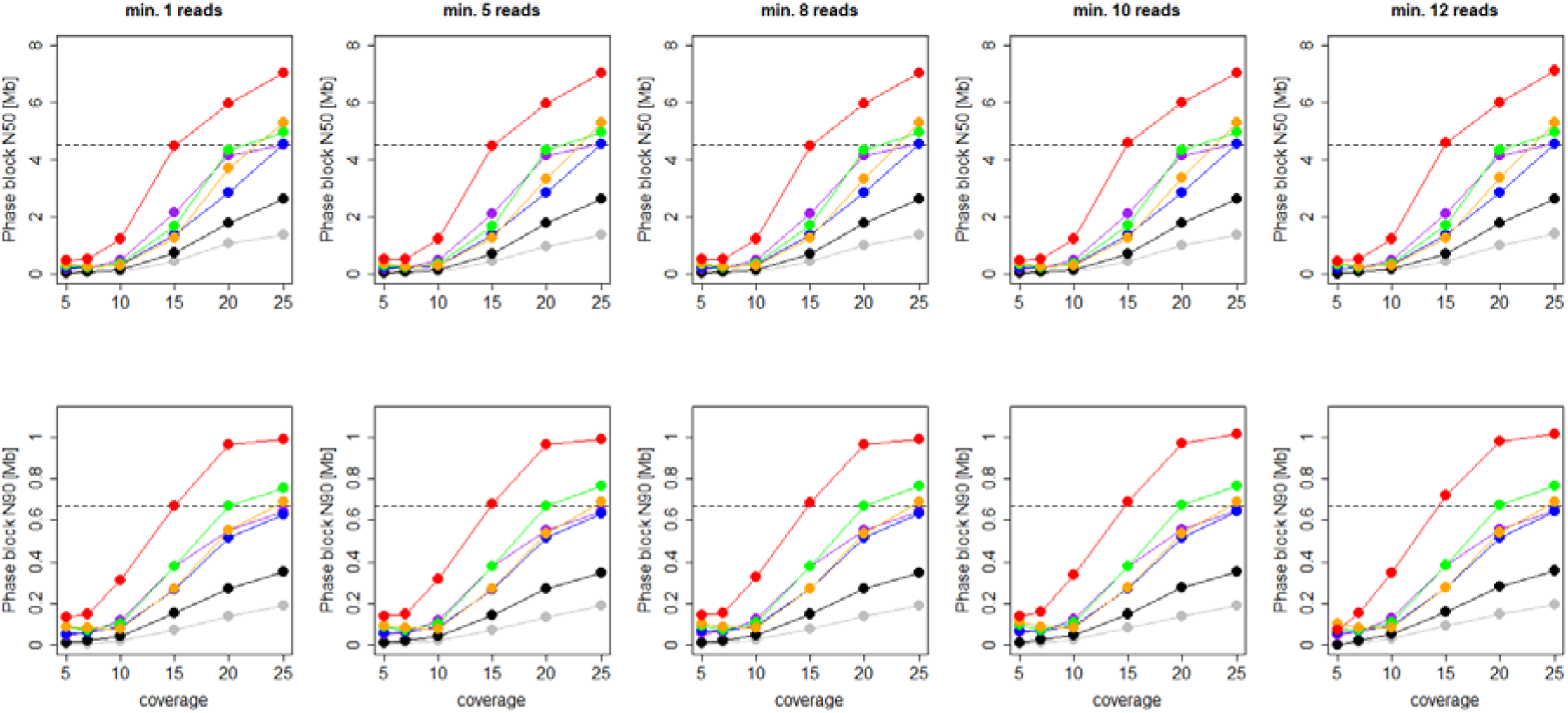
Phasing contiguity at different coverages and read-depth filtering. Different colors show values for different individuals/molecule lengths. Molecule lengths (kb): Red, 63.9; orange, 58.9; green, 47.8; blue, 37.3; purple, 33.1; black, 17.9; grey, 10.6. Broken lines show that similar contiguities can be reached with good-quality DNA (red) at lower coverage (15x) as with lower quality DNA (green, orange, blue, purple) at high coverage.

Contiguity in terms of phasing N50 (and N90) increased from about 10-530 kb (1-140 kb) at 5x coverage to 1,390-7,120 kb (190-1,020 kb) at 25x coverage (**Figure 1**). As shown by N50/N90 for full data, for input DNA >30 kb contiguity continues increasing at higher coverages (**Supplementary Figure S1**). At a given coverage, contiguity increased largely linearly with increasing input molecule length (**Supplementary Figure S2**). Filtering for a minimum read depth per SNP had no detectable effect on contiguity (p=0.92). The effect of molecule length increased with coverage, as indicated by their significant interaction (**Table 2**). Specifically, the gain in N50 increased from ca 8.5 kb per kb molecule length at 5x coverage to almost 90 kb per kb molecule length at 25x (**Supplementary Figure S3**). Molecule length seemed a better predictor of phasing contiguity at low and high than at intermediate coverages (see larger residuals for the latter in **Supplementary Figure S2**).

### Coverage of the genome by phase sets

The majority of the genome was covered by phase sets (that is sets of SNPs that are combined into haplotypes) provided adequate input DNA molecule length and a read-depth filtering of SNPs not exceeding average coverage (mean ± standard deviation: 87.1 ± 1.8 %, **Figure 2, Table 3**).

**Table 3.** Effect of coverage, molecule length, and filtering on coverage of genome by phase sets (multiple R^2^=0.66, 204 degrees of freedom).

|  | Estimate | t | p |
| --- | --- | --- | --- |
| Intercept | 74.586 | 82.042 | $< 10^{-3}$ |
| Coverage | 1.867 | 14.603 | $< 10^{-3}$ |
| Molecule Length | 0.246 | 4.994 | $< 10^{-3}$ |
| Filter | -1.968 | -8.372 | $< 10^{-3}$ |
| Coverage x Molecule Length | -0.028 | -4.100 | $< 10^{-3}$ |
| Coverage x Filter | 0.278 | 8.422 | $< 10^{-3}$ |

**Figure 2.**
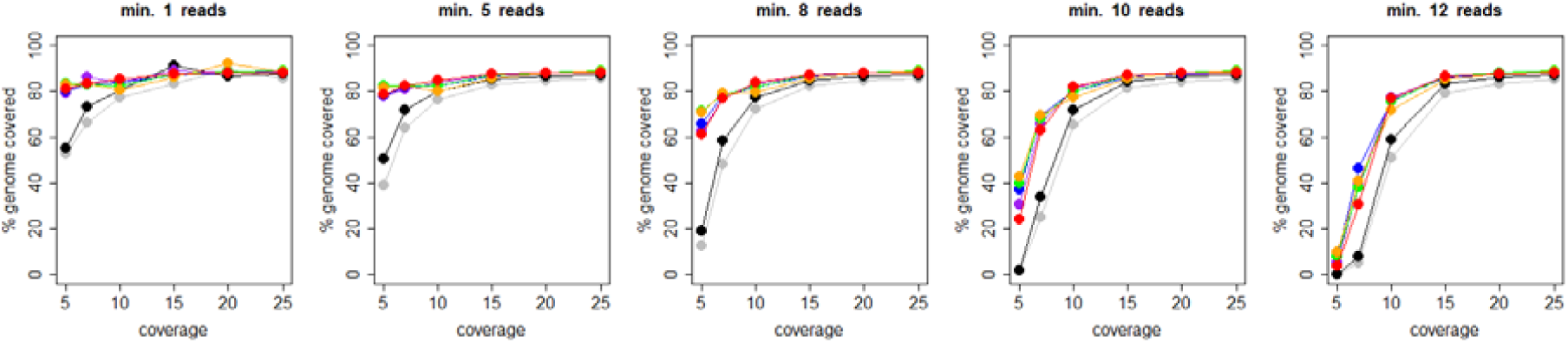
Proportion of the genome covered by phase sets at different coverages and read-depth filtering. Different colors show values for different individuals/molecule lengths. Molecule lengths (kb): Red, 63.9; orange, 58.9; green, 47.8; blue, 37.3; purple, 33.1; black, 17.9; grey, 10.6.

The proportion of the genome covered by phase sets increased with coverage and input molecule length and decreased with filtering stringencies that exceed sequencing coverage (**Figure 2, Table 3**). With input molecule lengths >30 kb, this proportion plateaued slightly below 90% for coverages starting at 15x, independent of filtering. Read-depth filters exceeding sequencing coverage drastically reduced the proportion of the genome covered by phase sets. This is expected, as such filtering simply removes most SNPs from the data set.

### Phasing and genotyping accuracy

Both phasing and genotyping accuracies strongly increased with increasing coverage and with adequate read-depth filtering (**Figure 3, Table 4**).Phasing accuracy increased from down to almost 60% at 5x coverage to above 99% starting at 15x coverage for input DNA with molecule lengths estimated >30 kb (**Figure 3, Table 4**). Samples with input DNA <20 kb still reached phasing accuracies >98%. Filtering for a minimum read-depth improved phasing accuracy significantly at low coverages (**Figure 3, Table 3**). Finally, at all but low coverages, phasing accuracy increased with increasing molecule length (**Supplementary Figure S4, Table 4**; at low coverage phasing accuracy was lower for long molecule lengths). Genotyping accuracy likewise strongly increased with increasing coverage (**Figure 3, Table 4**) and was further improved by adequate read-depth filters. Accuracies above 99% were reached starting at a coverage of 15x with a read-depth filter of at least 8. However, overfiltering (for higher than genome-wide average coverage) strongly decreased genotyping accuracy (**Figure 3**), as indicated by a significant interaction of filtering with coverage (**Table 4**). Finally, at coverages of 15x or higher, genotyping accuracy increased with increasing input molecule length, while the opposite effect was observed at lower coverages (**Supplementary Figure S5**).

**Table 4.** Effect of coverage, filtering, and molecule length on phasing accuracy (multiple R^2^=0.80, 204 degrees of freedom) and genotyping accuracy (multiple R^2^=0.85, 204 degrees of freedom).

|  | Phasing Accuracy |  |  | Genotyping Accuracy |  |  |
| --- | --- | --- | --- | --- | --- | --- |
|  | Estimate | T | p | Estimate | T | p |
| Intercept | -0.739 | -14.785 | $< 10^{-3}$ | -0.0743 | -21.378 | $< 10^{-3}$ |
| Coverage | 0.197 | 28.003 | 0.004 | 1.107 | 31.775 | $< 10^{-3}$ |
| Filtering | 0.037 | 2.891 | $< 10^{-3}$ | 0.284 | 8.168 | $< 10^{-3}$ |
| Molecule Length | 0.009 | 3.348 | $< 10^{-3}$ | 0.006 | 0.178 | 0.86 |
| Coverage x Filtering | -0.007 | -3.879 | $< 10^{-3}$ | 0.189 | 5.412 | $< 10^{-3}$ |
| Coverage x Molecule Length | 0.001 | 3.219 | 0.001 | 0.099 | 2.844 | 0.005 |

**Figure 3.**
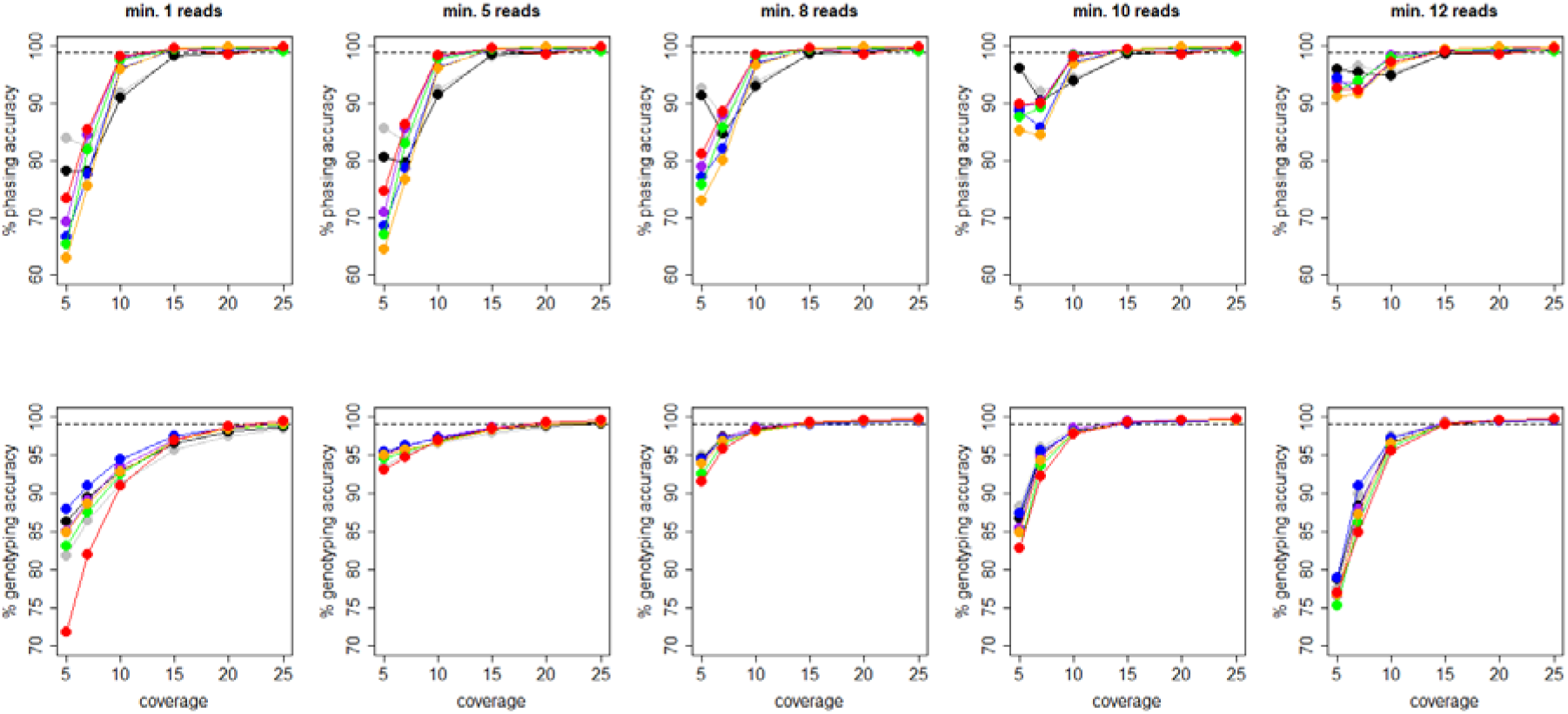
Phasing and genotyping accuracies at different coverages and read-depth filtering. Different colors show values for different individuals/molecule lengths. Molecule lengths (kb): Red, 63.9; orange, 58.9; green, 47.8; blue, 37.3; purple, 33.1; black, 17.9; grey, 10.6.

## Discussion

Our analyses suggest that phasing contiguity and accuracy, as well as genotyping accuracy of haplotype-resolved sequencing improve with increasing sequencing coverage and mean input DNA quality in terms of molecule length. Additional improvements of genotyping accuracy can be reached with adequate read-depth filtering. However, with adequate input DNA (> 30 kb), phasing contiguities at Mb-scale with high phasing accuracies in the same range as genotyping accuracy can be achieved already with a sequencing coverage of 15x. Thereby, the sequencing effort required to obtain haplotype-resolved genome resequencing data of high quality is in the same range as currently used in many large-scale population resequencing studies (e.g. Burri *et al.* 2015; Martin *et al.* 2013; Stankowski *et al.* 2019; Vijay *et al.* 2016), and thus affordable both financially and in terms of the amount of input DNA required. Our study thus reveals an affordable avenue towards haplotype-resolved genome resequencing data at population scale, even for organisms with gigabase-sized genomes like birds. In the following, we discuss the implications of our results and the recommendations deriving therefrom for the design of studies that aim to make use of haplotype-resolved sequencing.

### Towards population-scale haplotype-resolved sequencing: high coverage is silver, moderate coverage gold

Although sequencing at high yields highest phasing contiguity, moderate coverages around 15x open the avenue of high-quality phasing at populations scale.

According to our results, best phasing contiguities and accuracies for each sample are achieved at high sequencing coverage (**Figures 1, 3**). Phasing contiguity reached N50 of >7 Mb at 25x coverage, and above this coverage continued increasing up to almost 11 Mb, as shown when adding contiguities of full data with coverages >45x (**Supplementary Figure S1**). In contrast, phasing accuracy improved only marginally with coverages beyond 15x (**Figure 3**). From 15x coverage on it exceeded 99% and was thus in the same range as genotyping accuracy. In contrast to genotyping accuracy, which can be strongly improved by filtering for a minimum read-depth, gains in phasing accuracy through filtering were moderate and risked the cost of covering a reduced proportion of the genome with phasing information (**Figure 2**). Higher than 15x sequencing coverage thus predominantly affords improved phasing contiguities.

However, our prime interest was in establishing whether adequate phasing contiguities and accuracies can be achieved with affordable sequencing effort to enable haplotype-resolved sequencing at population-scale. Indeed, our results suggest that already with reasonable sequencing effort phasing contiguities adequate for most applications in molecular ecology can be reached. At a coverage of 15x of linked reads, which is similar to the coverage nowadays used in many population-scale short-read sequencing experiments (e.g. Burri *et al.* 2015; Martin *et al.* 2013; Stankowski *et al.* 2019; Vijay *et al.* 2016), phasing N50 reaches Mb-scale (**Figure 1**) and phasing accuracy exceeding 99%. For samples with input DNA >30 kb, the phasing accuracies we estimated at 15x and higher coverage are in the same range as observed in equivalent human data, where linked-read data consistently outperformed statistical phasing in all respects (Choi *et al.* 2018). Moreover, the phasing accuracies estimated here compare to values reported for human genomes sequenced with the same technology at 25x coverage (<0.4%, Porubsky *et al.* 2017). Not even with parent-offspring trios current state-of-the-art statistical phasing approaches reach as high phasing accuracy as we find at this moderate coverage (Al Bkhetan *et al.* 2019).

Thus already sequencing coverages of 15x should enable high-quality inferences of phase blocks relevant to many population genetic inferences in outbred natural populations. In cichlids, for instance, average ancestry tract lengths is at most 3 kb; the majority of ancestry tract is smaller than 20 kb, and only few are longer than 50 kb (Meier *et al.* 2017). In European sea bass, even in low-recombination regions migrant tracts are on average 50 kb long; longest migrant tracts measure up to 2 Mb (Duranton *et al.* 2018). The longer ancestry tract sizes in sea bass versus cichlids reflects the more recent admixture in this system that started only about 11,500 years ago as opposed to up to 100,000-200,000 years ago in cichlids. Still, even for the case of relatively recent admixture in sea bass, phasing contiguities achievable with 15x coverage should span the majority of haplotype tracts. However, in systems with more recent admixture with onsets a couple of hundred years ago, such as between North American canids, ancestry tracts can be considerably longer, averages reaching several Mb (up to >9 Mb) (vonHoldt *et al.* 2011). In species with such a recent history of admixture, phasing contiguities achieved with 15x coverage indeed may be shorter than relevant haplotype tract lengths. However, additional statistical phasing of phase blocks with read-aware software that treats phase blocks as reads, such as WhatsHap (Martin *et al.* 2016), may extend the phasing beyond relevant contiguities even for such recent cases of admixture.

In conclusion, our results suggest that with the same sequencing efforts as used in standard resequencing studies linked-read sequencing is able to provide valuable phasing information, thus paving the way for affordable haplotype-resolved genome resequencing at population scale.

### Issues with tissues: Phasing contiguity requires high-quality input DNA

Our results suggest that requirements for high phasing contiguities start with the choice of tissues for DNA extraction, or latest in the wet lab with adequate high molecular weight (HMW) DNA extraction methods. The physical size over which linked-read sequencing can provide haplotype information is in first line limited by the size of input DNA fragments. In line with this expectation, we observed highest phasing contiguities for DNA of high quality, that is with long input DNA molecules >30 kb; DNA of poor quality (<20 kb) did not yield good phasing contiguity. Moreover, for DNA of good quality the same phasing contiguity can be reached at low to moderate coverage as with poor DNA at high coverage (broken line in **Figure 1**). Still, at intermediate coverages, the correlation of phasing contiguity with input molecule length is poorer, likely due to the stochasticity with which fragments are sequenced at these coverages. At these coverages the chance of fragments that are part of a long molecule to not be sequenced increases, and long molecules may be broken up into shorter phase sets. Nevertheless, best phasing contiguities require high-quality DNA. Therefore, tissue preservation and DNA extraction methods that yield HMW DNA help keeping sequencing costs affordable and may be key to take haplotype-resolved genome sequencing to population scale.

The importance of good DNA quality to obtain high phasing contiguities and keep sequencing costs at a minimum implies that measures to obtain the best possible source of DNA must be taken starting from tissue collection on. Systematic studies on the choice and preservation of tissue samples are yet lacking. Although our study was not designed to infer best practices for tissue preservation and DNA extraction for linked-read sequencing, our results suggest that DNA quality depends on the tissue of origin. We extracted DNA exclusively with a magnetic bead-based HMW DNA protocol, and neither tissue age nor preservation buffer (in all cases ethanol) are related to differences in DNA quality. The observed differences in molecule lengths thus likely reflect issues with the type of tissues used for DNA extraction. Indeed, the most notable characteristic of the two data sets with input molecule lengths <20 kb was the tissue of origin: in both cases this was muscle as opposed to blood for the other samples (**Table 1**). Therefore, to preserve the best possible source for HMW bird DNA, we recommend that bird tissue collections include samples of blood (red blood cells are nucleated in birds). For museum-based cryo-collections, which often exclusively sample solid tissues from body cavities but not blood, this implies that blood be collected prior to sacrificing birds. After tissue collection, measures to protect DNA from damage are required, starting with the preservation in the lab/field and ending with DNA extraction. The suitability of different preservation buffers remains to be investigated. However, we recommend that samples be cooled at the best possible, and that DNA extraction makes use of methods dedicated to the isolation of HMW DNA (Klingström *et al.* 2018).

While genomic studies have been limited by sequencing technology to date, sequencing now starts being limited by the quality of input DNA. We thus recommend that research in the wild today starts collecting tissues with qualities for tomorrow.

## Acknowledgements

We are grateful to the Yale Peabody museum for providing tissue of a male *O. melanoleuca* (101348) and to the Museum of Vertebrate Zoology of Harvard University for providing tissue of a male *O. pleschanka* (349924). We thank Ana Gomes for extracting HWM DNA, and acknowledge support from the National Genomics Infrastructure in Stockholm funded by the Science for Life Laboratory, the Knut and Alice Wallenberg Foundation and the Swedish Research Council, and the SNIC/Uppsala Multidisciplinary Center for Advanced Computational Science for assistance with massively parallel sequencing and access to the UPPMAX computational infrastructure. Further computation was performed at the High-Performance Computing Cluster EVE, a joint effort of the Helmholtz Centre for Environmental Research (UFZ) and the German Centre for Integrative Biodiversity Research (iDiv) Halle-Jena-Leipzig. We thank the administration and support staff of EVE: Thomas Schnicke and Ben Langenberg (UFZ), and Christian Krause (iDiv). This research was supported by a Science for Life Laboratory Swedish Biodiversity Program grant (2015-R14) to AS and by a German Research Foundation (DFG) research grant (BU3456/3-1) to RB.

## Author contributions

RB designed the research with input from DL. DL, RR, RB, and HSch performed data analysis. JG and PE performed library preparation and genome sequencing. RAO performed genome assembly. MS, AS, JTG, and HShi contributed materials. RB wrote the paper with input from all authors.

## Supplementary Figures

**Supplementary Figure S1.**
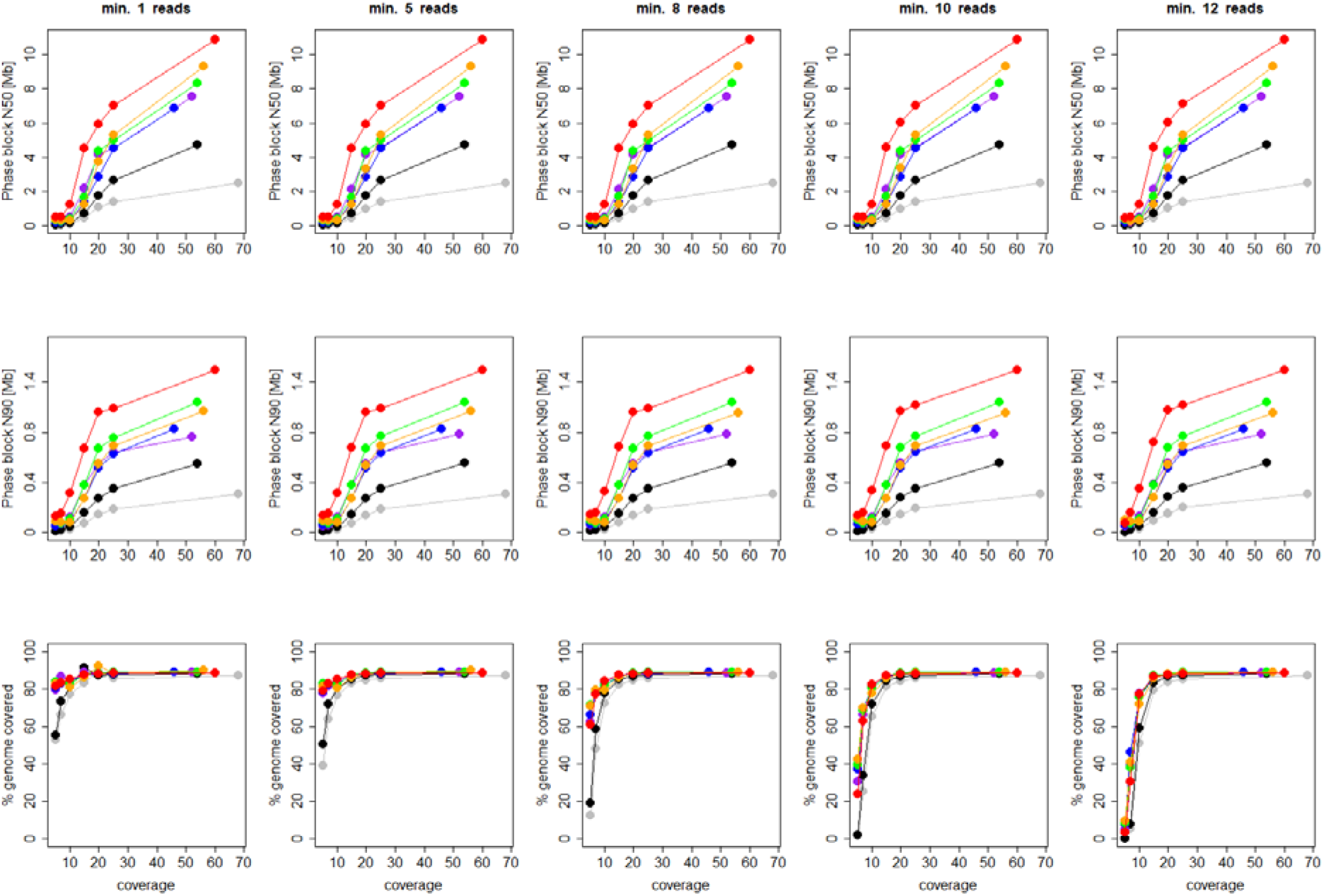
Contiguity and genome coverage by phase sets including full, not downsampled data. Different colors show values for different individuals/molecule lengths. Molecule lengths (kb): Red, 63.9; orange, 58.9; green, 47.8; blue, 37.3; purple, 33.1; black, 17.9; grey, 10.6.

**Supplementary Figure S2.**
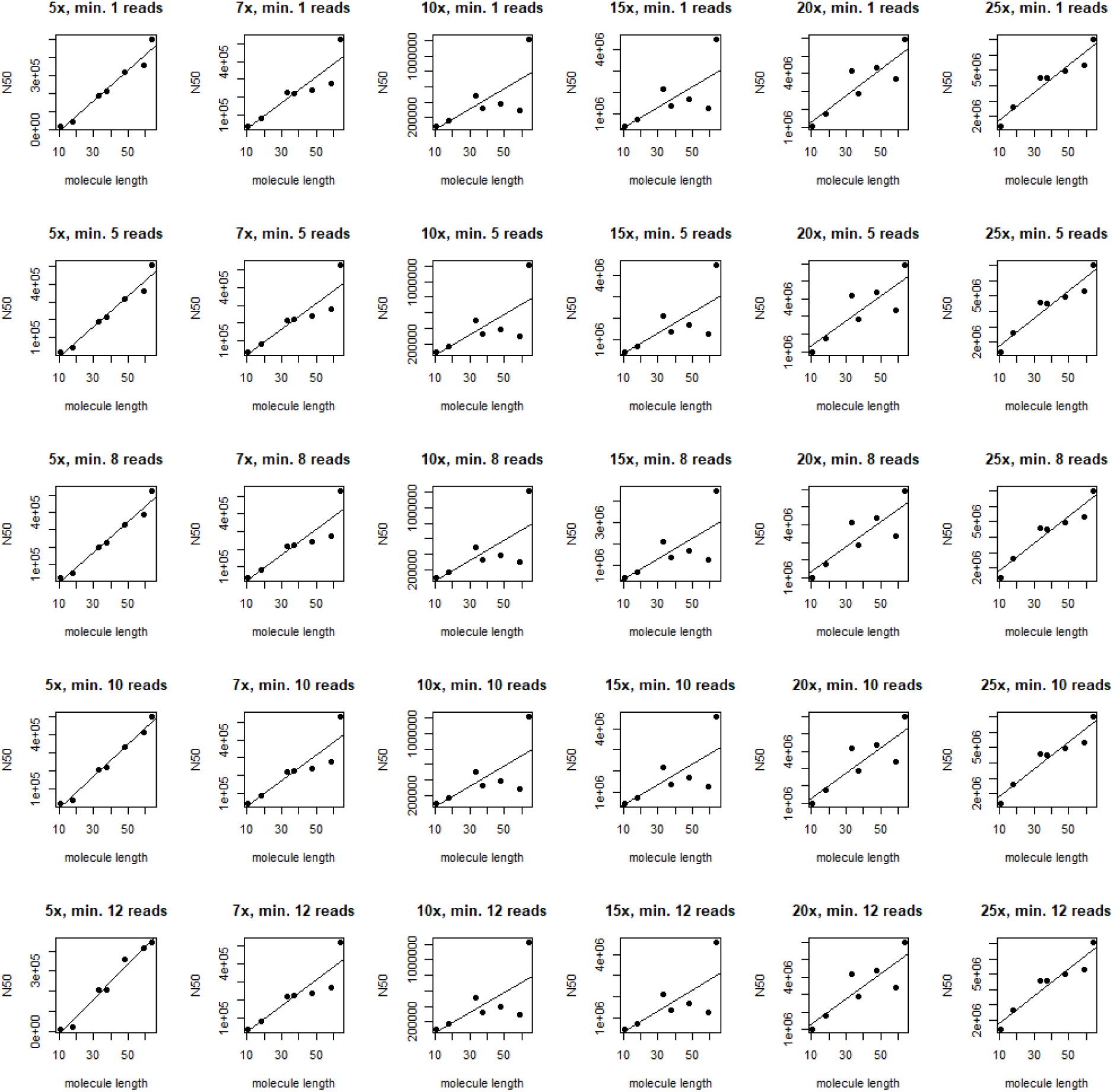
Effect of molecule length on contiguity separately for each coverage and filter. Different colors show values for different individuals/molecule lengths. Molecule lengths (kb): Red, 63.9; orange, 58.9; green, 47.8; blue, 37.3; purple, 33.1; black, 17.9; grey, 10.6.

**Supplementary Figure S3.**
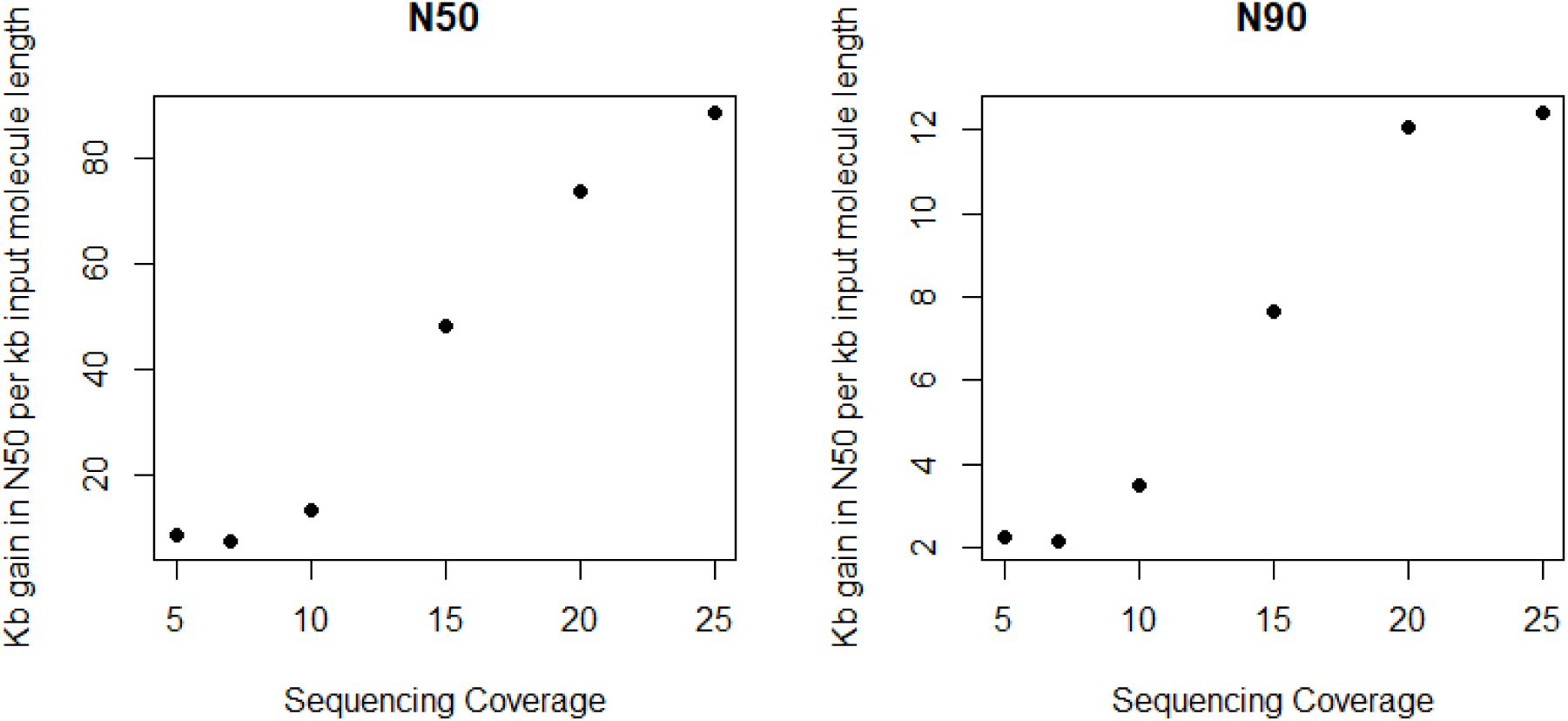
Gain in contiguity through increased input molecule lengths at different coverages.

**Supplementary Figure S4.**
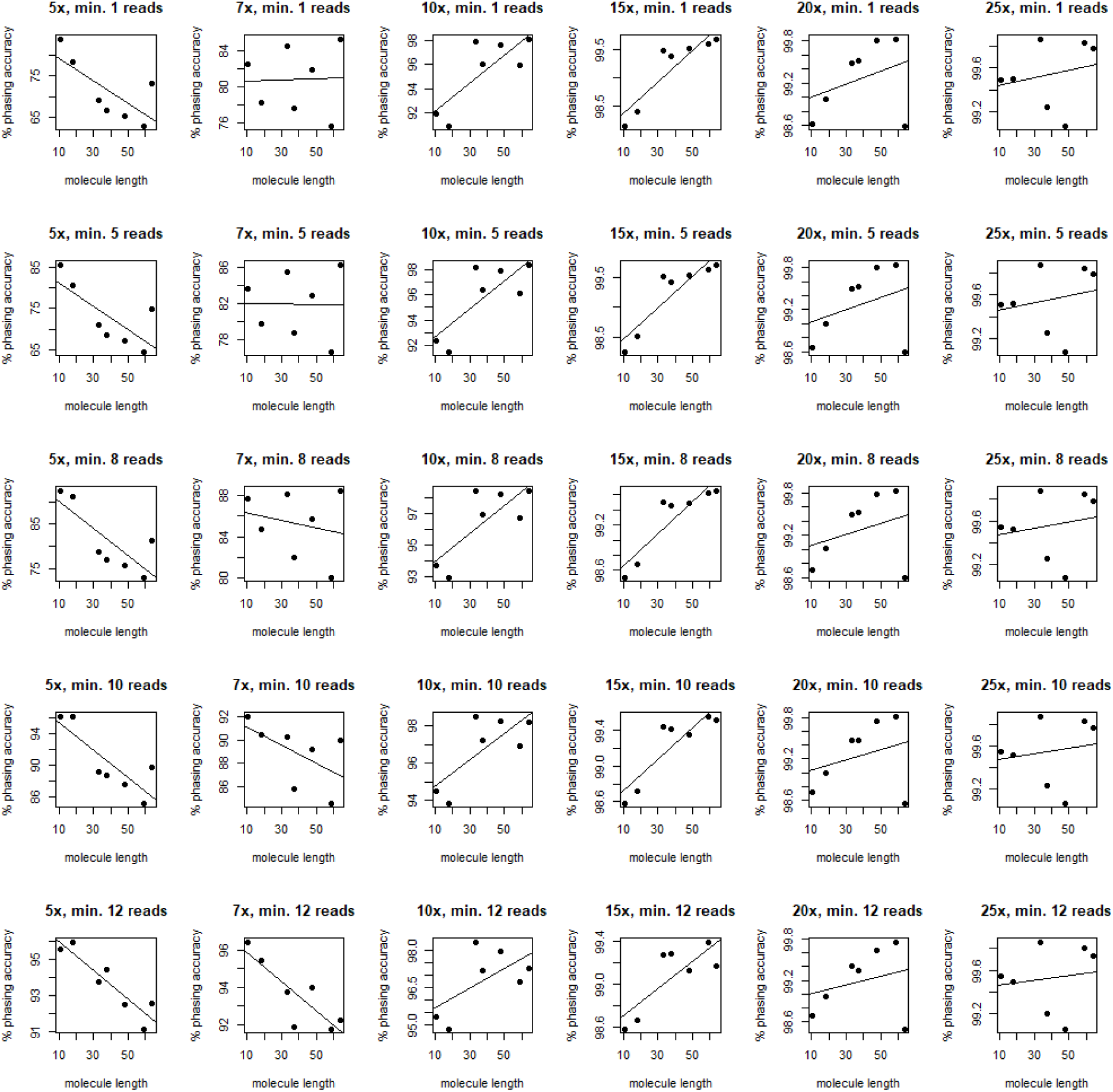
Effect of molecule length on phasing accuracy separately for each coverage and filter. Different colors show values for different individuals/molecule lengths. Molecule lengths (kb): Red, 63.9; orange, 58.9; green, 47.8; blue, 37.3; purple, 33.1; black, 17.9; grey, 10.6.

**Supplementary Figure S5.**
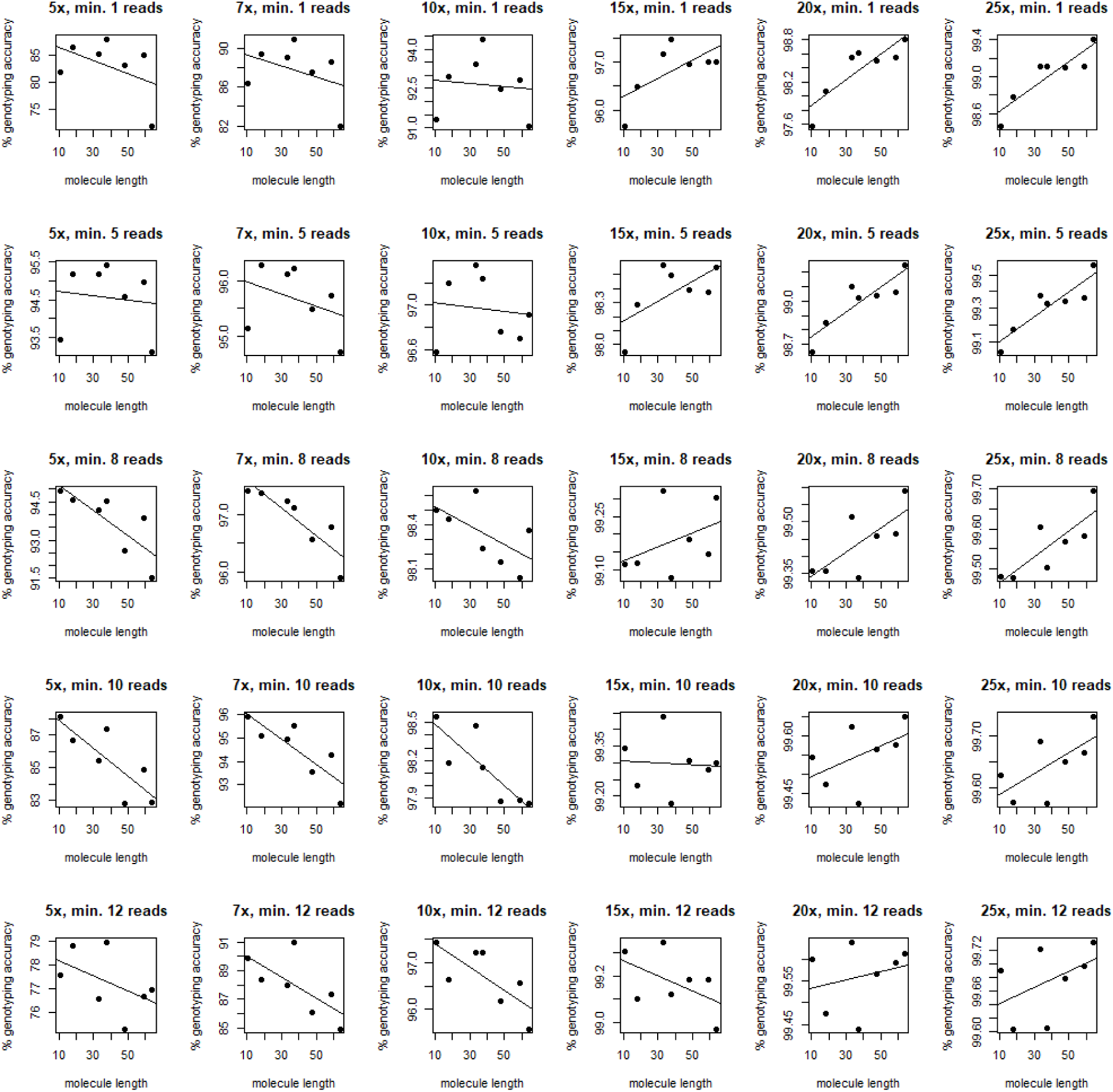
Effect of molecule length on genotyping accuracy separately for each coverage and filter. Different colors show values for different individuals/molecule lengths. Molecule lengths (kb): Red, 63.9; orange, 58.9; green, 47.8; blue, 37.3; purple, 33.1; black, 17.9; grey, 10.6.

